# Host Diet Influences Lethal and Sublethal Responses of Hosts to Amphibian Pathogen Exposure

**DOI:** 10.1101/290239

**Authors:** Samantha L. Rumschlag, Michelle D. Boone

**Affiliations:** Department of Biology, Miami University, Oxford OH

**Keywords:** host-pathogen interactions, wildlife disease, host quality, *Batrachochytrium dendrobatidis*, chytridiomycosis, anurans

## Abstract

The severity of the impacts of pathogens on hosts may be driven by environmental factors like resource availability that create tradeoffs on energetic demands for immune responses and basal metabolic activity within the host. These responses can vary among species from sublethal to lethal effects, which can have consequences for the host population trajectories within a community. Chytridiomycosis, caused by the pathogen *Batrachochytrium dendrobatidis* (or Bd), has been associated with global amphibian population declines. However, it also occurs in populations without appearing to cause mass mortality; the effect of Bd in these situations is not well understood and environmental factors like food abundance that impact host conditions could play an important role in the magnitude of the pathogen’s impact. In the present study, we exposed American toad (*Anaxyrus americanus*), northern leopard frog (*Lithobates pipiens*), and Blanchard’s cricket frog (*Acris blanchardi*) metamorphs to Bd and then reared them in the terrestrial habitat under low or high food environments. We found additive effects of Bd and reduced food abundance on host growth and survival that varied according to species. For instance, Bd-induced reductions in American toad survival were greater under low food conditions compared to high food conditions but survival of northern leopard frogs and Blanchard’s cricket frogs was not affected by Bd. For northern leopard frogs and Blanchard’s cricket frogs, low food abundance resulted in the lower growth rates under Bd exposure compared to high food abundance. Additionally, we developed stage-structured population models for American toads to assess if reduced survival of metamorphs exposed to Bd under conditions of low and high food abundance could influence population trajectories; models indicated that Bd exposure would reduce annual population growth rates by 14% under conditions of high food abundance and 21% under conditions of low food abundance. Our results suggest that environmental conditions that influence resource availability for species that are sensitive or tolerant to Bd may increase the negative effects of Bd on host growth and survival, which could have important implications for how populations and communities with infected members respond over time.

## Introduction

Interspecific variation in the responses of hosts to infectious pathogens can drive host-pathogen interactions with consequences for changes in host population dynamics and the transmission of pathogens throughout a community (Sait et al. 1994, Boots et al. 2003, Fenton and Pedersen 2005). When exposed to a pathogen, hosts can exhibit a wide range of responses. For instance, mortalities rates of hosts may vary among species, driving differences in population trajectories (Fenton and Pedersen 2005). In addition, sublethal effects of pathogens may have important consequences for populations and communities, and may influence the spread of disease to other hosts. Sublethal effects of pathogens, which often receive less attention in disease ecology, can reduce fecundity, increase the time to development, and may destabilize populations (Sait et al. 1994, Boots et al. 2003). Understanding the variation in both the lethal and sublethal responses of hosts across species is critical for predicting and preparing for outbreaks of infectious diseases in wildlife, yet currently we have limited information to predict patterns observed in nature. Energetic resources available for hosts that can aid in the defense against infection and disease development could be a key factor influencing disease dynamics, which may provide predictive power in forecasting impacts of infectious diseases on host populations. Under conditions in which few food resources are available, such as droughts or increases in temperature, the energy devoted to immune responses compared to basal metabolic activities (e.g. growth, development, behavior, reproduction) may be less (Blaustein et al. 2012). Tradeoffs on energetic demands between immune responses and basal metabolic activities that occur for hosts of small size may contribute to the development of sublethal or lethal effects of pathogens in hosts that are otherwise unaffected or only suffer sublethal effects.

The effects of chytridiomycosis, a disease caused by the fungal pathogen *Batrachochytrium dendrobatidis* (hereafter, Bd), may be magnified under conditions of low food resource availability and induce lethal and sublethal effects on amphibian hosts. Bd has been called the most damaging infectious disease on vertebrates in modern history (Murray et al. 2011); the pathogen can infect upwards of 500 species of amphibians from all continents on which amphibians exist (Olson et al. 2013). Exposure of amphibian hosts to Bd can result in sublethal effects on host growth (Bielby et al. 2015, Caseltine et al. 2016), mortality (Kleinhenz et al. 2012, Wise et al. 2014), and has been linked to population declines of amphibians around the world (Berger et al. 1998, Muths et al. 2003, Lips et al. 2006). We understand less about the disease ecology of chytridiomycosis in temperate regions like the midwestern United States compared to places like Central and South America and Australia where mass mortality events have been sudden and widespread. This gap in understanding limits our perception of the range of host-pathogen interactions in this system that can structure communities and drive population dynamics.

Because mounting an immune response is an energetically costly process (Lochmiller and Deerenberg 2000), larger hosts with greater energetic reserves may be better able to sustain pathogenic exposures without the risk of impacts on survival and growth. Host conditions and fitness is predicted by body size across taxa (Dobson 1992, Bachman and Widemo 1999, Shine et al. 2001). For instance in amphibians, larger body size is associated with earlier time to first reproduction, increased fecundity, and increased overwinter survival (Smith 1987, Scott et al. 2007, Earl and Whiteman 2015). Larger individuals, which have superior host condition, may have lower risks associated with pathogen exposure because of a better ability to mount energetically costly immune responses.

The impact of host size on amphibian response to pathogen exposures may be significant at metamorphosis, which is a critical time for development of the immune system of amphibians (Rollins-Smith 1998). Metamorphosis in amphibians is a time of complete reorganization of organ systems which may leave metamorphs temporarily vulnerable to pathogens (Rollins-Smith 1998). Exposure of hosts to pathogens near metamorphosis may increase the risk of lethal and sublethal impacts on hosts, which may be compounded by poor host conditions driven by availability of food resources.

The objective of the present study is to determine the influence of host condition, as measured by body size, on the effect of exposure to Bd on three temperate species: the American toad (*Anaxyrus americanus*), the northern leopard frog (*Lithobates pipiens*), and the Blanchard’s cricket frog (*Acris blanchardi*). All three species can be infected with Bd in the field (Longcore et al. 2007, Goodman and Ararso 2012, Richards-Hrdlicka et al. 2013), and American toads and northern leopard frogs suffer effects on survival or growth of Bd when exposed (Ortiz-Santaliestra et al. 2013, Wise et al. 2014, Caseltine et al. 2016). Additionally, northern leopard frogs and cricket frogs are declining in parts of their ranges, which could be linked to Bd (Hecnar and M’Closkey 1996, Rorabaugh 2005, Voordouw et al. 2010). We hypothesized low food abundance, which determines size and condition of hosts, would increase the likelihood of negative effects of Bd exposure in all three anuran species.

## MATERIALS AND METHODS

### Animal Collection and Care

Six northern leopard frog (*Lithobates pipiens*) egg masses were collected on 2 April 2014 from Talawanda High School Pond (39°29’16”N, 84°43’42”W). Seventeen partial egg strings of American toad (*Anaxyrus americanus*) were collected on 12 April and 18 April 2014 from Rush Run Wildlife Area (39°34’59”N, 84°37’4”W). Eggs from four amplexed Blanchard’s cricket frog (*Acris blanchardi*) pairs were collected on 7 June and 9 June 2014 from a pond at Miami University’s Ecology Research Center (39°31’43”N, 84°43’25”W). Eggs were held at 17°C on a 12:12 h light-dark cycle until they reached the free-swimming stage (Gosner stage 25 [Gosner 1960]). Tadpoles were fed ground TetraMin Tropical fish flakes (Tetra Holding) ad libitum until they were transferred to outdoor mesocosm ponds at Miami University’s Ecology Research Center (Oxford, OH, USA). Thirty northern leopard frog, American toad, or Blanchard’s cricket frog tadpoles, were added to mesocosms on 17 April, 22 April 2014, and 19 June 2014, respectively. Each mesocosm pond contained 1000 L water, 1 kg leaf litter, plankton inoculates, and tadpoles of a single frog species (American toad, northern leopard frogs, or Blanchard’s cricket frogs), and a fiberglass screen lid. We reared American toad tadpoles in 17 mesocosm ponds, northern leopard frogs in 5 mesocosm ponds, and Blanchard’s cricket frogs in 10 mesocosm ponds. The number of mesocosm ponds varied because American toads were reared for multiple studies and because we anticipated variation in rates of metamorphosis among species. Tadpoles were reared in mesocosm ponds through metamorphosis when they were transferred to the laboratory. A subsample of metamorphs of each species reared in mesocosms was used in the terrestrial portion of the current study.

We housed frogs individually within terraria in plastic shoebox containers (41.3 x 17.8 x 15.6 cm) that contained layers of pea gravel (∼1.5 cm) and topsoil (∼2.5 cm), a small dish for water, and an upturned dish for cover. Frogs were held at 22°C on a 14:10 h light-dark cycle. Species were reared for different lengths of time because of the phenology of metamorphosis differed among species; however, all three studies were concluded at the same date on (2 September 2014). This resulted in the terrestrial, post-metamorphosis portion of the experiment lasting 10 weeks for American toads, 8 weeks for northern leopard frogs, and 4 weeks for Blanchard’s cricket frogs.

### Experimental Design

In the terrestrial portion of the experiment, we manipulated exposure to Bd (present, absent) and food abundance (low, high) with 20 replicates of each treatment for American toads, northern leopard frogs, and Blanchard’s cricket frogs for a total of 80 experimental units per species. Treatments were assigned randomly to individual frogs within a species so that the experimental unit was the individual frog.

We exposed post-metamorphic anurans to Bd for 12 hr on 26 June 2014 for American toads, 9 July 2014 for northern leopard frogs, 6 August 2014 for Blanchard’s cricket frogs. To expose anurans to Bd, we placed individuals in ventilated plastic petri dishes with 8 mL dechlorinated water and 1 mL of the assigned treatment solution (see below). After 12 hr, frogs were returned to their assigned terrarium. We cultured Bd (isolate JEL 213 isolated from *Rana muscosa* in the Sierra Nevada [USA], obtained from J. Longcore, University of Maine, Orno, ME) on 1% tryptone agar plates using standard protocols (Longcore et al. 1999). Bd zoospores were harvested with 3 mL dechlorinated water. For Bd-absent treatments, we added dechlorinated water to 1% tryptone agar plates without Bd cultures. After 30 min, we collected the water from the plates into two solutions, one containing Bd zoospores and the other that was absent of Bd. We calculated zoospore concentrations using a hemocytometer, and diluted the Bd zoospore solution with dechlorinated water so that all hosts were exposed to 1.25×10^6^ zoospores/mL. To test for initial Bd infection, two weeks after Bd exposure, we euthanized ten anurans of each species that had been exposed to Bd (five fed on a high food diet, five fed on a low food diet) using a 1% solution of MS-222 (tricaine methanesulfonate), stored them in ethanol, and sent swabs of their bodies to the Amphibian Disease Lab at the San Diego Zoo for qPCR testing for the presence of Bd.

Frogs were fed calcium-dusted crickets in the terrestrial portion of the experiment three times per week. We manipulated feeding regime at two levels: low and high. Anurans in the low-feeding treatment received amount of crickets approximately equal to 2% of their mean body mass before Bd exposure. Each week that the frogs in the low-food abundance consumed all the crickets presented, we increased the amount of food by either one or two crickets or a cricket size class (0.3175 cm, 0.635 cm, and 1.27 cm). The high-food abundance was always three times as many crickets as the low treatment. High-food abundances were essentially *ad libitum* because uneaten crickets were not removed from containers. We observed survival daily, and individual frogs were weighed weekly to measure growth.

### Statistical Analyses

We tested for the effects of food abundance, Bd exposure, and the interaction of these treatments on American toad and Blanchard’s cricket frog survival using logistic regression. All northern leopard frogs survived the course of the experiment indicating that food abundance, Bd exposure, and the interaction of these treatments did not impact survival. We used repeated-measures analysis of variance (ANOVA) to determine the effects of food abundance, Bd exposure, and the interaction of these treatments on log-transformed mass of northern leopard frogs and cricket frogs over the course of the experiment. We used repeated-measures ANOVA to test for the effects of food abundance and Bd exposures on log-transformed mass of American toads; the interaction of food abundance and Bd exposure was not included in the statistical model because of low survival of American toads, which led to missing cells. To assess the effects of treatments on size of individuals over the course of the experiment, we used an ANOVA to test for the effects of food abundance, Bd exposure, and the interaction of these treatments on change in mass (final mass – initial mass [before Bd exposure]) of northern leopard frogs and cricket frogs. We used an ANOVA to test for the effects of food abundance and Bd exposure on change in mass (final mass – initial mass [before Bd exposure]) of American toads; the interaction of food abundance and Bd exposure was not included in the statistical model because of low survival of American toads that led to missing cells. All analyses were completed using SAS 9.2 (SAS Institute, Inc., Cary, North Carolina). ANOVAs were constructed using generalized linear models (PROC GLM) with a Gaussian distribution, and results were evaluated using Type III error with α = 0.05. Logistic regressions (PROC LOGISTIC) were built with a binary distribution and a logit link function, and results were evaluated using Type III analyses of effects with α = 0.05.

### Population Model

To consider the influence of Bd and food abundance on host population growth, we built stage-structured Lefkovitch (Caswell 2000) annual projection matrices representing female populations of American toads with a birth pulse (Biek et al. 2002). Only American toads were modeled because they were the only species for which we found significant effects of Bd on toad survival. We modeled American toads under four conditions: no exposure to Bd with high food abundance, no exposure to Bd with low food abundance, exposure to Bd with high food abundance, and exposure to Bd with low food abundance. Our models were composed of three life stages: pre-juvenile (embryo, larva, and overwintering metamorph), juvenile, and reproductive adult (Biek et al. 2002). The projection matrix that representing an American toad population under high food abundance and no exposure to Bd is:

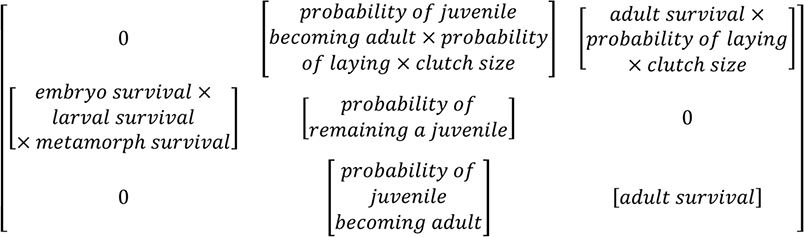

To represent the effects of reduced food abundance and Bd exposure, metamorph survival was reduced according to our experimental results. For instance, to model low food abundance with no Bd exposure, metamorph survival was reduced by 17% compared to metamorph survival with high food abundance and no Bd exposure. Similarly, to represent Bd exposure under conditions of high food abundance, metamorph survival was reduced by 72%, and to show Bd exposure under conditions of low food abundance, we reduced metamorph survival by 94%.

Matrix elements consisting of vital rates of American toads or related species were taken from the scientific literature (Table 1). Mean embryo survival rates and standard distributions are based off observations of hatching success of American toad egg masses (Miller 1909, Harris et al. 2000, Allran and Karasov 2001, McDaniel et al. 2004). Mean larval survival and standard deviations are from experimental observations of Woodhouse’s toad (*Anaxyrus woodhousii*) (Boone et al. 2004) and long-term field observations of wood frogs (*Lithobates sylvantica)* (Berven 1990). Mean metamorph survival to spring emergence and standard deviation are from survival rates of American toad metamorphs in terrestrial enclosures where groups of 10 recently metamorphosed toads were overwintered in 2×2 m terrestrial enclosures (Distel and Boone 2010). The mean juvenile survival rate and standard deviation is from an estimate of juvenile survival of boreal toads (*Anaxyrus boreas*; Biek et al. 2002).

**Table 1.**
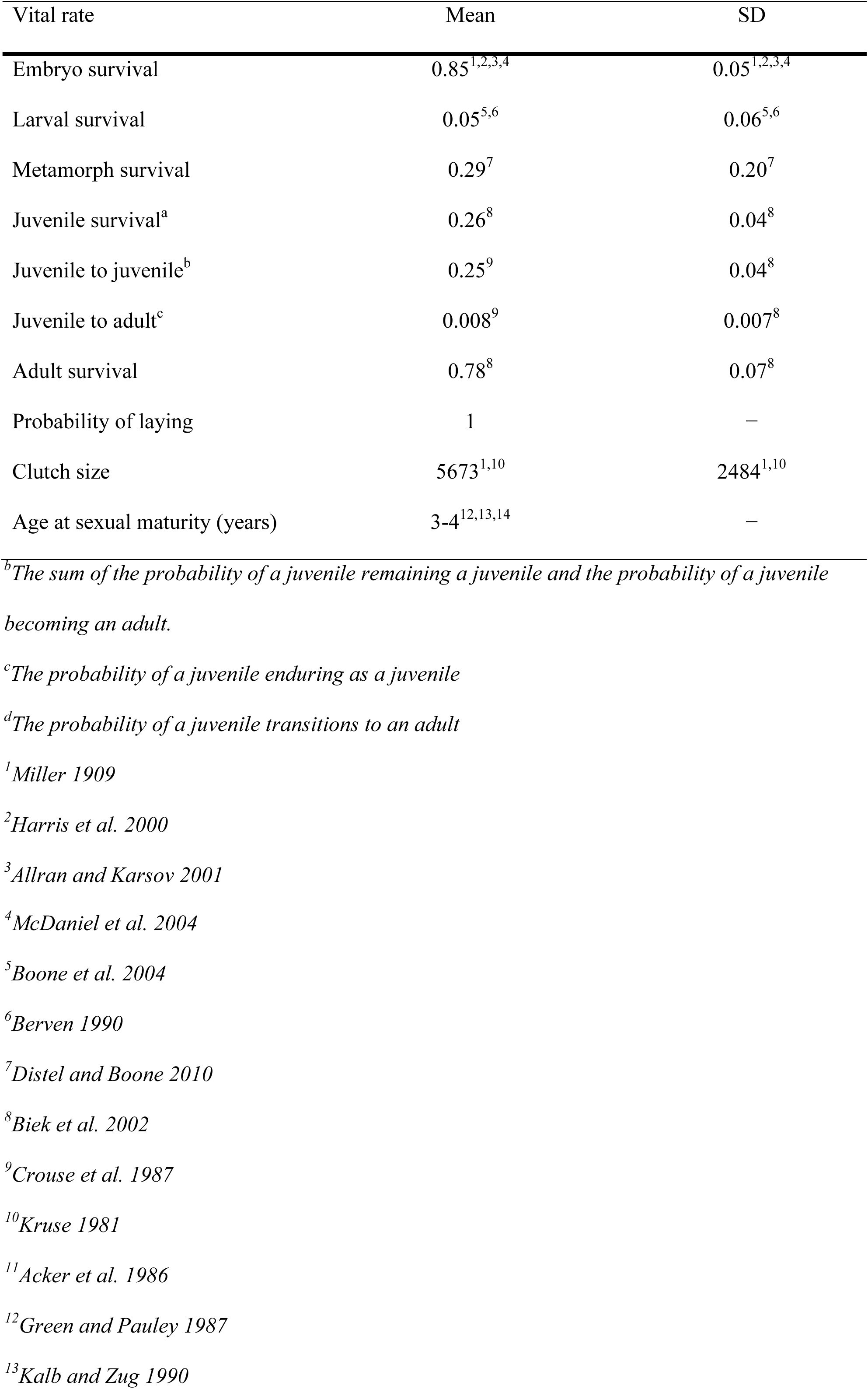
Mean vital rates, matrix elements, and corresponding standard deviations (SD) of stage-structured Lefkovitch projection matrices representing an American toad population. Vital rates and matrix elements represent annual transitions for females with the exception of embryo, larval, and metamorph survival, which combined represent a single year.

| Vital rate | Mean | SD |
| --- | --- | --- |
| Embryo survival | 0.85<sup>1,2,3,4</sup> | 0.05<sup>1,2,3,4</sup> |
| Larval survival | 0.05<sup>5,6</sup> | 0.06<sup>5,6</sup> |
| Metamorph survival | 0.29<sup>7</sup> | 0.20<sup>7</sup> |
| Juvenile survival<sup>a</sup> | 0.26<sup>8</sup> | 0.04<sup>8</sup> |
| Juvenile to juvenile<sup>b</sup> | 0.25<sup>9</sup> | 0.04<sup>8</sup> |
| Juvenile to adult<sup>c</sup> | 0.008<sup>9</sup> | 0.007<sup>8</sup> |
| Adult survival | 0.78<sup>8</sup> | 0.07<sup>8</sup> |
| Probability of laying | 1 | — |
| Clutch size | 5673<sup>1,10</sup> | 2484<sup>1,10</sup> |
| Age at sexual maturity (years) | 3-4<sup>12,13,14</sup> | — |

<sup>b</sup>The sum of the probability of a juvenile remaining a juvenile and the probability of a juvenile becoming an adult.
<sup>c</sup>The probability of a juvenile enduring as a juvenile
<sup>d</sup>The probability of a juvenile transitions to an adult
<sup>1</sup>Miller 1909
<sup>2</sup>Harris et al. 2000
<sup>3</sup>Allran and Karsov 2001
<sup>4</sup>*McDaniel et al. 2004* <sup>5</sup>*Boone et al. 2004* <sup>6</sup>*Berven 1990* <sup>7</sup>*Distel and Boone 2010* <sup>8</sup>*Biek et al. 2002* <sup>9</sup>*Crouse et al. 1987* <sup>10</sup>*Kruse 1981* <sup>11</sup>*Acker et al. 1986* <sup>12</sup>*Green and Pauley 1987* <sup>13</sup>*Kalb and Zug 1990*

To estimate the mean probability of a juvenile remaining a juvenile, we used the following formula: P_1_ = ((1-p_i_^di-1^) * p_i_) / (1-p_i_^di^) where p_i_ is the annual probability of survival for a juvenile and d_i_ is the number of years spent as a juvenile (Crouse et al. 1987). In our model, the female American toads reach sexual maturity in 3-4 years (Acker and Krehbiel 1986, Green and Pauley 1987, Kalb and Zug 1990). We estimated the mean probability of a juvenile remaining a juvenile by taking the average of P_1_ when d_i_ equals three and P_1_ when d_i_ equals four. The formula used for the mean probability of a juvenile becoming an adult is P_2_ = (p_i_^di^ * (1-p_i_)) / (1-p_i_^di^) (Crouse et al. 1987). The mean probability of a juvenile becoming an adult is the average of P_2_ when d_i_ equals three and P_2_ when d_i_ equals four.

The mean adult survival rate and standard deviation used in the model is from an estimate of adult survival of boreal toads (*Anaxyrus boreas*; Biek et al. 2002). We assumed a probability of females laying a clutch of 1 with a standard deviation of 0. Mean clutch size and standard deviations are based off of counts of American toad clutches (Miller 1909, Kruse 1981). The models characterize female American toads, so clutch size is halved under the assumption of a 1:1 sex ratio.

We calculated λ, the finite rate of increase of population growth, at stable age distribution for 2000 matrices that we generated by drawing randomly from a log-normal distribution of clutch sizes and β-distributions for all other vital rates. These distributions were built with 2000 observations using means and standard deviations in Table 1 as in Biek et al. (2002). We used sensitivity and elasticity analyses on mean vital rates to determine how small changes in each vital rate would influence λ when all other vital rates are held constant (De Kroon et al. 2000). Modeling exercises were completed in R version 3.2.1. with code adapted from Stevens (2010).

## Results

Results from qPCR analyses revealed high infection prevalence two weeks after exposure to Bd: American toads (high food: 1.0, low food: 1.0), northern leopard frogs (high food: 0.6, low food: 0.8), cricket frogs (high food: 1, low food: 0.75) (species [high food abundance: proportion infected, low food abundance: proportion infected]).

### Manipulating Food Resources and Bd Exposure

Northern leopard frog (1.00 ± 0) and Blanchard’s cricket frog (0.98 ± 0.018) survival was not influenced by treatments or their interactions (mean survival ± standard error) (Table 2). Food abundance and Bd exposure, but not the interaction of the two treatments influenced American toad survival over 70 days (Table 2). While American toads exposed to Bd experienced reduced survival, American toads fed more food had a greater chance of survival in both the Bd-exposed and unexposed treatments (Figure 1).

**Figure 1.**
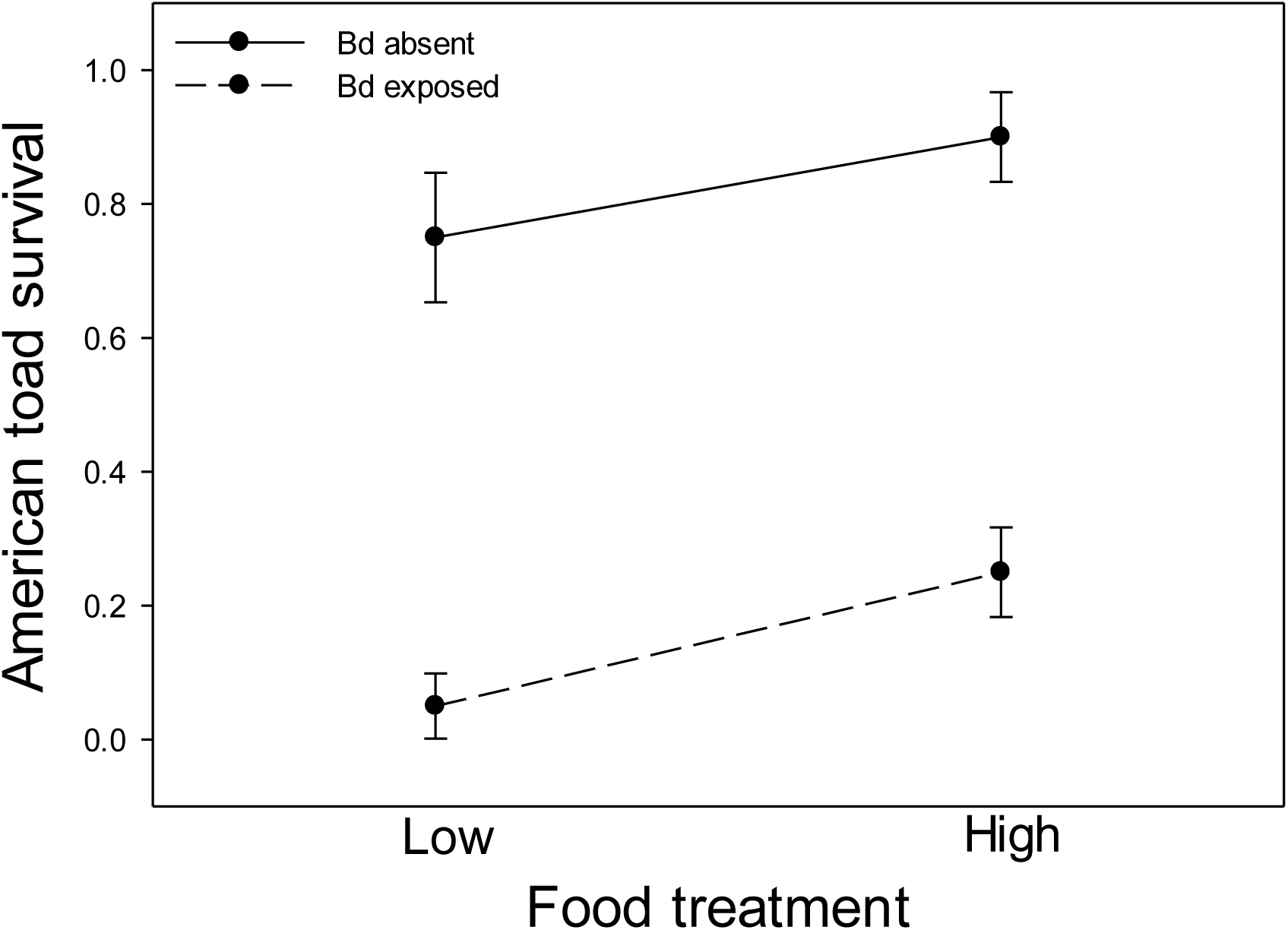
Survival over 70 days of American toads (*Anaxyrus americanus*) that were given different amount of food (low, high) and Bd treatments (absent, exposed). Plotted values are means ± binomial SE.

**Table 2.**
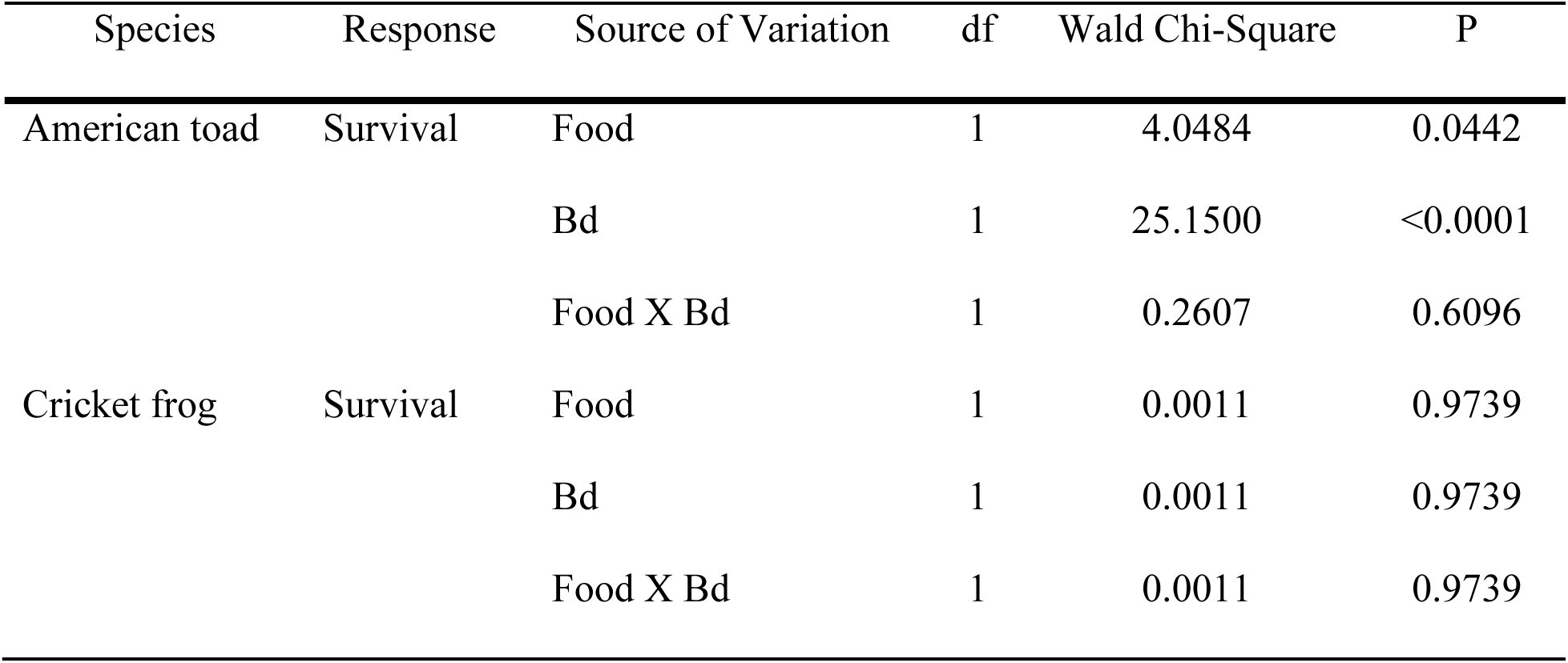
Summary of logistic regressions assessing the impact of food (low, high) and Bd (absent, exposed) treatments and their interaction on survival of American toads (*Anaxyrus americanus*) and northern cricket fog (*Acris blanchardi*). All northern leopard frogs (*Lithobates pipiens*) survived.

High food abundance significantly increased the mass of American toads, northern leopard frogs, and Blanchard’s cricket frogs over time (Table 3; Figure 2). Northern leopard frog mass over time was also influenced by Bd exposure (Table 3) with lower mean mass of northern leopard frogs exposed to Bd compared to the controls (Figure 2b). There was no effect of Bd or the interaction of Bd and food abundance on mass of American toads or cricket frogs over the course of the experiments (Table 3).

**Figure 2.**
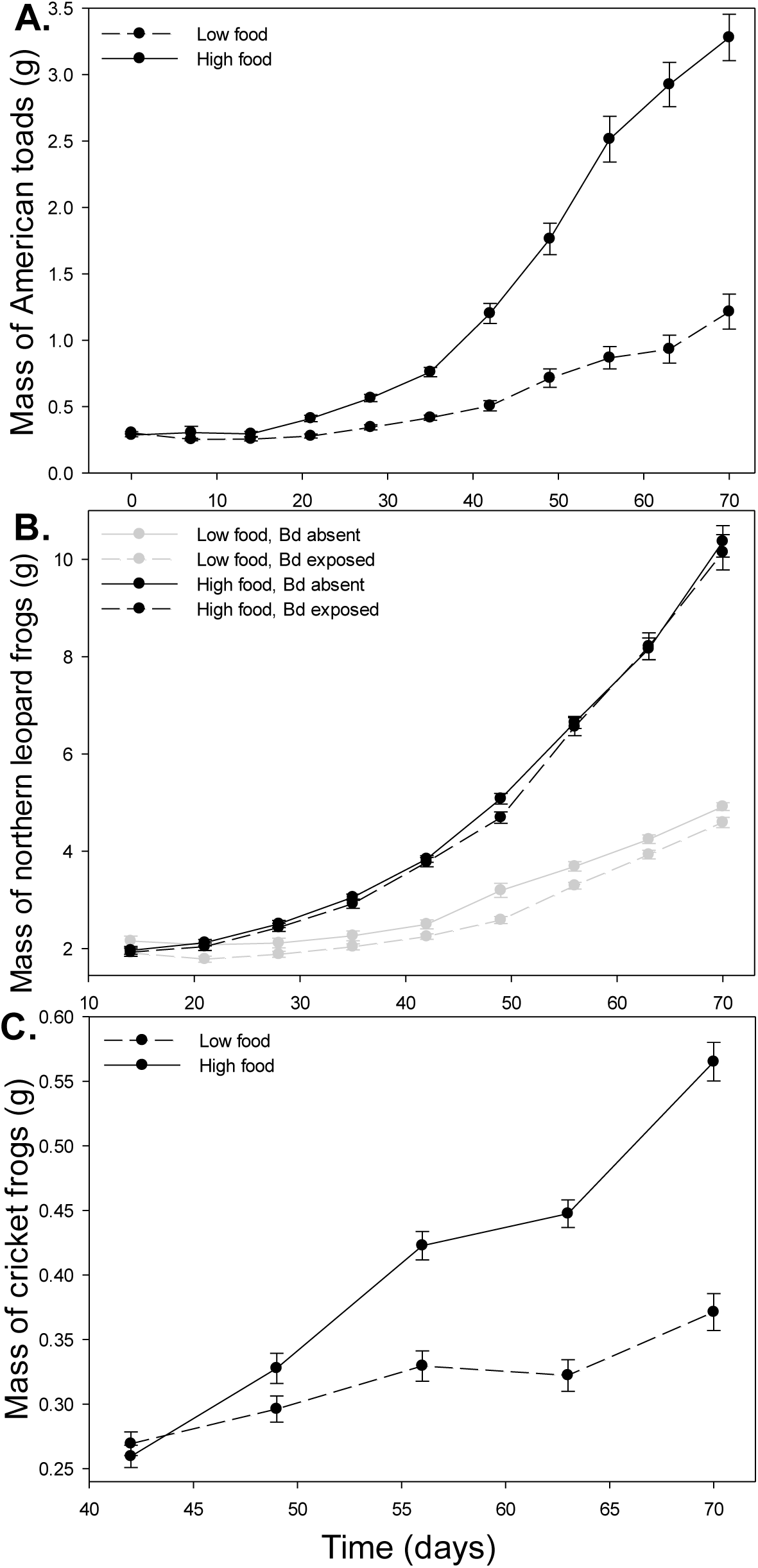
Mass over the course of the experiments of A) American toads (*Anaxyrus americanus*) in response to low and high food abundances, B) northern leopard frogs (*Lithobates pipiens*) in response to food (low, high) and Bd treatments (absent, exposed), and C) Blanchard’s cricket frogs (*Acris blanchardi*) in response to low and high food abundances. Plotted values are means ± SE.

**Table 3.**
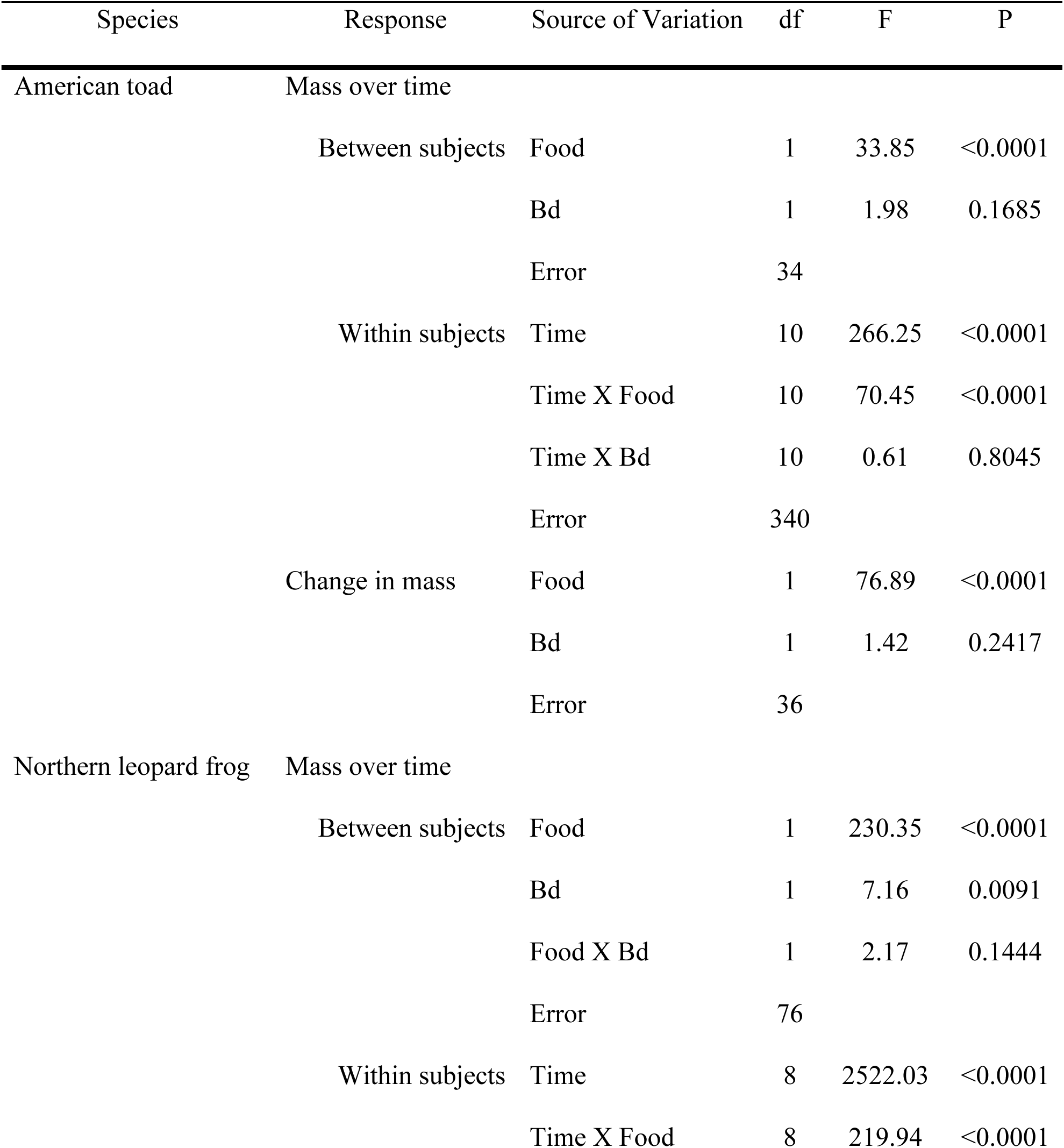
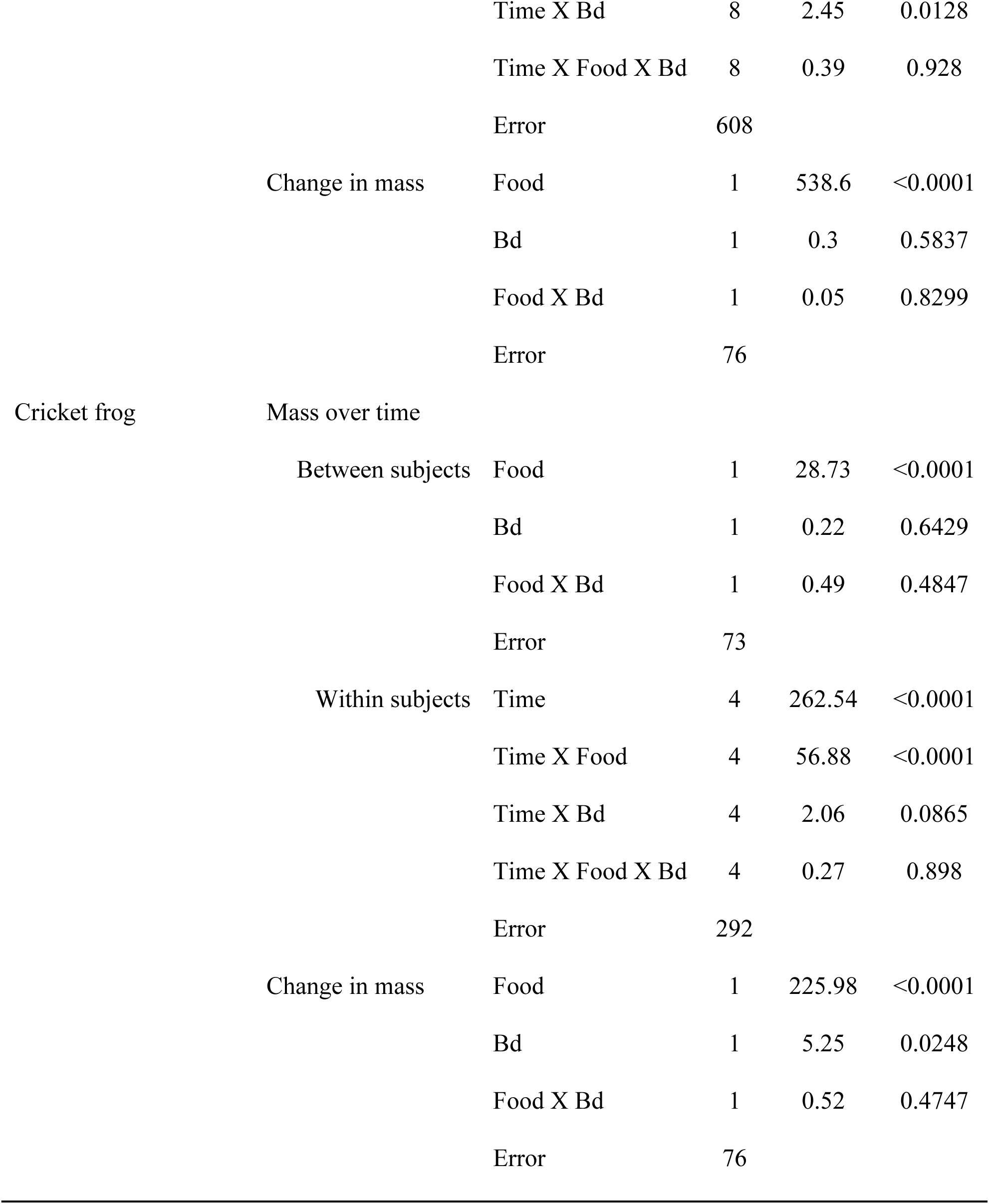
Summary of repeated measures ANOVAs and ANOVAs of the impacts of food (low, high) and Bd (absent, exposed) treatments and their interaction on mass over time and change in mass (final mass - initial mass) on American toads (*Anaxyrus americanus*), northern leopard frogs (*Lithobates pipiens*), and cricket frogs (*Acris blanchardi*).

Similarly, high food abundance significantly increased the amount of mass gained in American toads, northern leopard frogs, and Blanchard’s cricket frogs (Table 3; Figure 3). Bd, but not the interaction of food abundance and Bd, impacted change in mass of cricket frogs (Table 3); Blanchard’s cricket frogs exposed to Bd gained less mass over the course of the experiment compared the control (Figure 3c).

**Figure 3.**
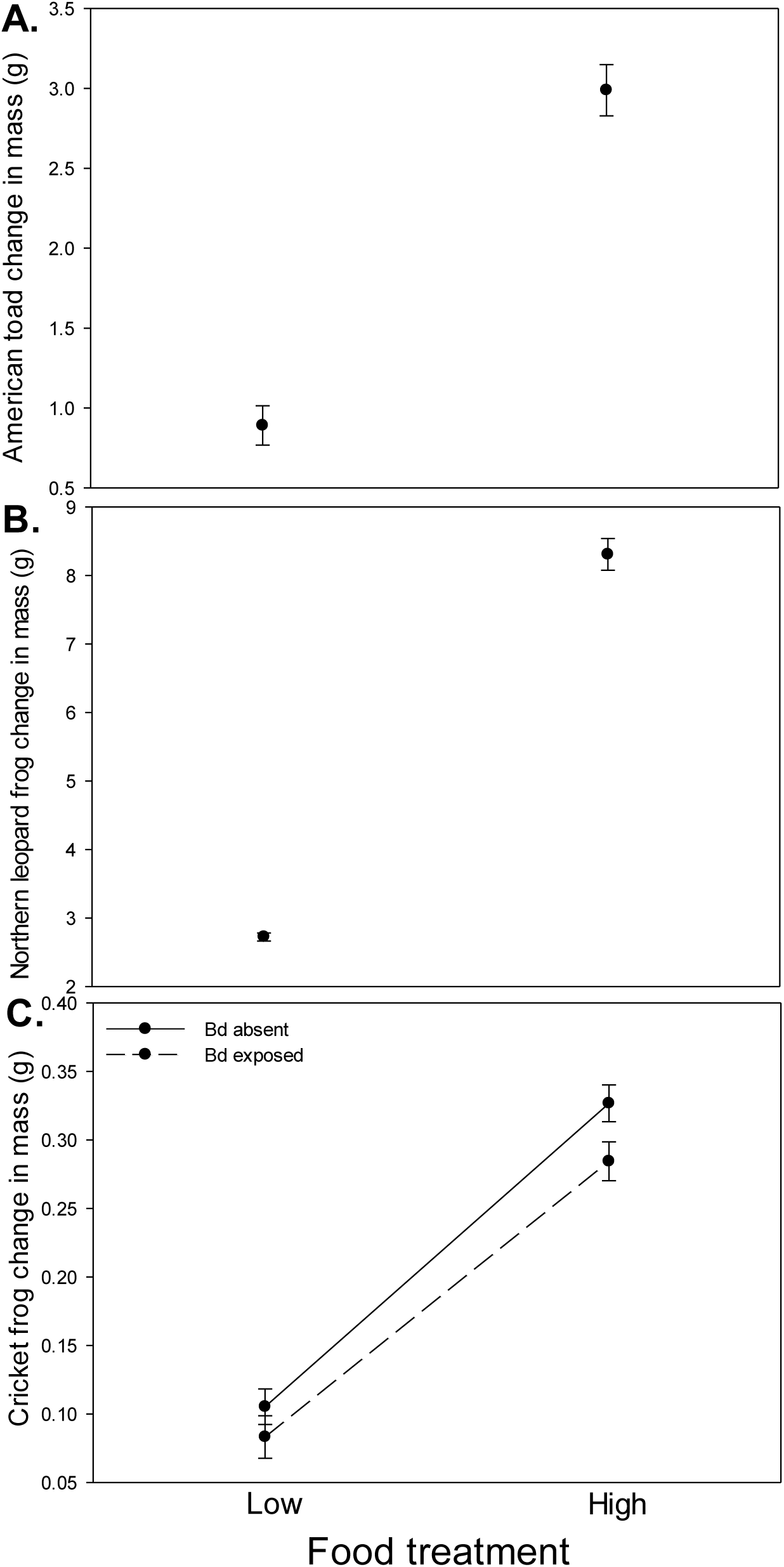
Change in mass (final mass – initial mass) of A) American toads (*Anaxyrus americanus*) in response to low and high food abundances, B) northern leopard frogs (*Lithobates pipiens*) in response to low and high food abundances, and C) Blanchard’s cricket frogs (*Acris blanchardi*) in response to low and high food abundances. Plotted values are means ± SE.

### Population Model

Bd exposure, represented as decreases in metamorph survival, decreased the finite rate of population growth λ over low and high food abundance, with the lowest population growth occurring under conditions of low food abundance and Bd exposure (Figure 4). The mean estimate of λ was 1.01 and 0.98 in the models that represented no Bd exposure under conditions high and low food abundance respectively. When we represented Bd exposure by reducing metamorph survival by 72% and 94% under conditions of high and low food abundance, the mean estimate of λ decreased by 14% and 21%, respectively, relative to the model of no Bd exposure and high food abundance (Figure 4).

**Figure 4.**
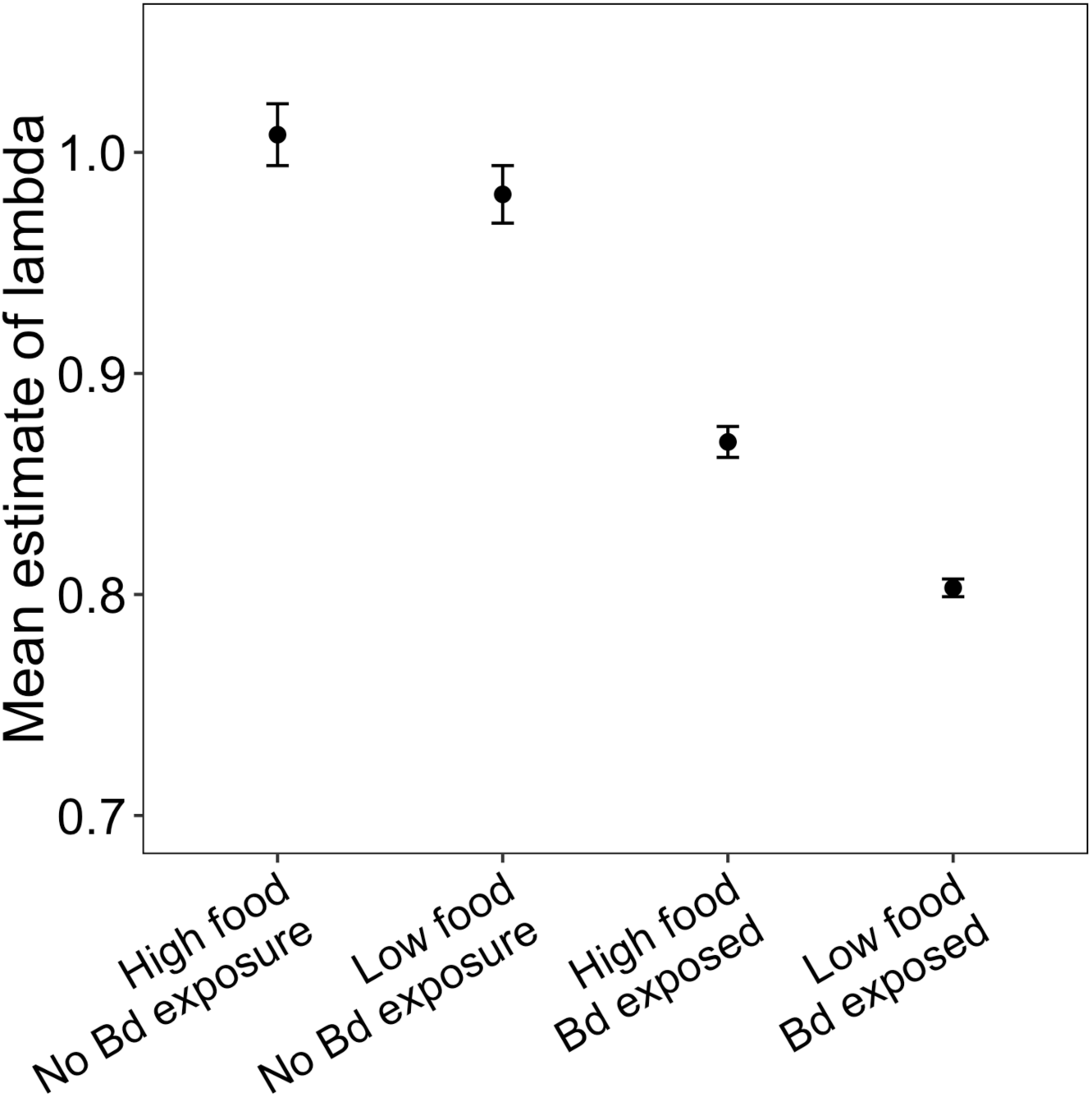
Mean estimates of λ with 95% confidence intervals for American toad populations that represent the influence of food abundance (low, high) and Bd exposure (absent, exposed) on metamorph survival to the juvenile stage.

Sensitivity analysis on the annual projection matrix representing populations of American toads not exposed to Bd and under conditions of high food abundance, showed that λ was most sensitive to changes in survival from the juvenile to adult stage followed closely by the pre-juvenile (embryo, larva, metamorph) to juvenile stage relative to the other matrix elements (Table 4). For the three other annual projection matrices representing no Bd exposure under low food abundance and Bd exposure under low or high food abundance, sensitivity analyses support that changes in the transition probability of pre-juveniles to juveniles would cause the biggest changes in λ (Table 4). Across the four projection matrices, elasticity analyses showed λ was most elastic to changes in adult survival. Small proportional changes in this transition element relative to the other elements would have the greatest impact on λ.

**Table 4.**
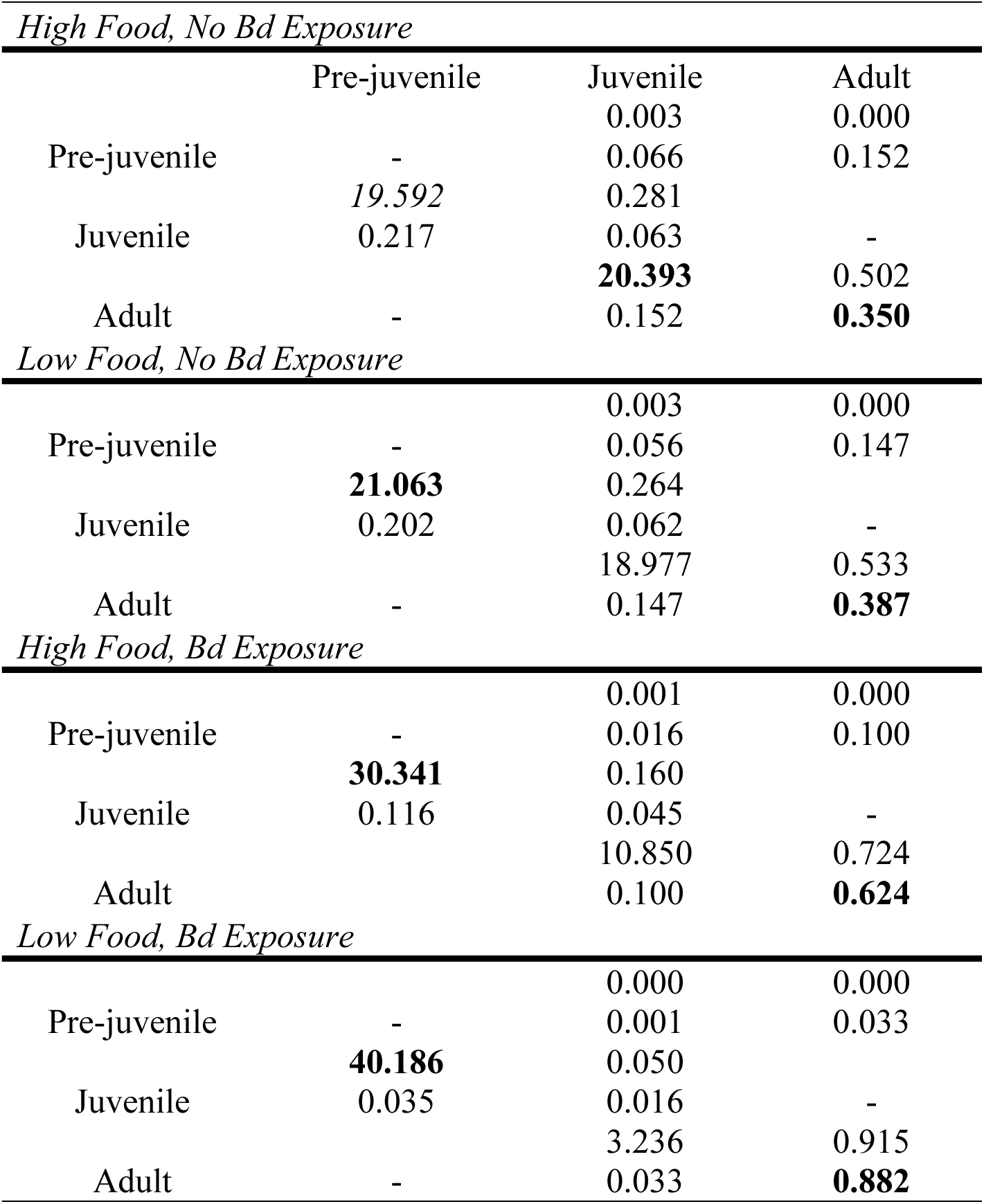
Sensitivity and elasticity values for projection matrices representing the following conditions: high food and no Bd exposure, low food and no Bd exposure, high food and Bd exposure, low food and Bd exposure. The first numbers listed in a column refers to sensitivity values, and the second refers to elasticity values. The greatest sensitivity and elasticity values for each model are in bold.

## Discussion

Change in resource availability represents perhaps one of the most widespread sources of environmental variation (e.g. Fretwell 1972). Increased frequency and intensity of droughts and extreme flow events (Barnett et al. 2005, Milly et al. 2005) associated with climate change could alter resource availability in vulnerable freshwater systems, which might result in alterations of host-pathogen interactions. Our study provides evidence demonstrating that food availability can mediate the lethal and sublethal impacts of pathogens on hosts. We found additive effects of low food abundance with Bd exposure on host growth and survival; individuals that were exposed to Bd and low food had the worst outcomes of any treatments.

Exposure to pathogens and fluctuations in food availability are both fundamental drivers of the health of organisms, which have consequences for population dynamics of organisms (Morin 2009). Our results show that both Bd exposure and low food availability negatively impact the growth and survival of amphibians across species. For American toads, low food abundance and Bd exposure lead to reductions in survival, with Bd exposure accounting for a larger effect compared to differences in food abundance. For northern leopard frogs and Blanchard’s cricket frogs, low food abundance and Bd exposure resulted in reductions in growth, with differences in food abundance accounting for a greater effect compared to Bd exposure. Together, these results suggest that American toads are more susceptible to the effects of Bd in the post-metamorphic life stage, while northern leopard frogs and cricket frogs may be more susceptible to low food availability.

While the effects of pathogens on hosts can vary across species, there may be environmental conditions that increase these negative effects on host health for both tolerant and susceptible hosts. Our results demonstrate that Bd exposure and low food abundance most negatively affected individual performance when applied in combination. The lowest rates of survival for American toads and growth for northern leopard frogs and Blanchard’s cricket frogs resulted when Bd exposure and low food abundance were combined. These results are similar to other studies that have found an increased likelihood of mortality in anurans of small body sizes exposed to Bd (Carey et al. 2006, Garner et al. 2009). The effects of Bd on hosts of poor condition may be driven by their reduced ability to mount an effective immune response and may be a common phenomenon. Mounting an immune response is an energetically costly processes (Lochmiller and Deerenberg 2000) and energetic tradeoffs for hosts of small body sizes between host growth and survival versus immune response may exist (Blaustein et al. 2012). Across species, larger hosts were better able to sustain exposure of Bd as evidenced by reduced impacts of Bd on survival or growth. Likely, these animals were better able to mount an immune response because of increased availability of energetic reserves, decreasing the impacts of Bd exposure. Our results support that host body size may be a predictor for the ability of hosts to respond to infectious pathogens and suggests that environmental conditions that reduce host condition like increased competition, drought, and pond drying anticipated with global climate change could increase the consequences of pathogens for hosts. These environmental conditions that result in reductions in prey availability may increase the effects of pathogenic exposures through changes in host condition with implications for host-pathogen interactions in this system.

While the impacts of Bd on temperate populations of amphibians in the Midwestern United States are generally unknown, because more research focus is given to areas in which mass mortality events have been sudden and widespread, our results indicate that increased mortality rates and decreased growth of hosts caused by pathogenic exposures under suboptimal conditions may influence population trajectories for these species within a community. American toads, northern leopard frogs, and Blanchard’s cricket frogs can use the same ponds for breeding and be present at ponds concurrently; combined with our research results, we propose that Bd may impact amphibian communities in subtle, but potentially dramatic ways over time through impacts on reduced fitness and recruitment. American toads may be especially vulnerable to competition by these more tolerant species under conditions of low food abundance. Population models of American toads show that decreases in metamorph survival may lead to negative impacts on population growth rates via reduced recruitment with the lowest population growth rates occurring when toads are exposed to Bd under conditions of low food abundance.

While Bd exposure did not influence survival of northern leopard frogs and Blanchard’s cricket frogs, we are not suggesting that their population trajectories may be unaffected by Bd in natural populations. Northern leopard frogs and Blanchard’s cricket frogs experienced reduced growth as a result of Bd exposure, which can lead to later time to first reproduction, decreased fecundity, and decreased overwinter survival (Smith 1987, Scott et al. 2007, Earl and Whiteman 2015). Northern leopard frogs and cricket frogs are declining in parts of their ranges, and sublethal impacts of Bd exposure could contribute to these enigmatic declines (Hecnar and M’Closkey 1996, Rorabaugh 2005, Voordouw et al. 2010). The impact of infectious pathogens in the absence of mass mortality is understudied in disease ecology, even though infectious pathogens can reduce fecundity, increase time to development, and are predicted to destabilize populations (Sait et al. 1994, Boots et al. 2003).

Resource availability can be a major driver of community interactions across ecosystems (Morin 2009). Our results provide evidence that food abundance can additively influence the effects of pathogen exposures on lethal and sublethal impacts of pathogens on hosts. We support that decreased growth and survival of hosts exposed to pathogens under conditions of low food availability may have important ramifications for host population dynamics increasing the potential for host population declines via reduced recruitment, fecundity, and overwinter survival. Monitoring amphibian communities for population-level consequences may provide insights into causes of enigmatic declines for species like the northern leopard frog and Blanchard’s cricket frog, which may appear to be tolerant to Bd infection because of the lack of mass mortality events of these species in the Midwest but suffer subtle effects that impact populations. Accurate predictions of environmental disturbance that change resource availability should consider changes to host-pathogen systems if we are to design effective management strategies to protect vulnerable populations.

## Acknowledgments

We are grateful to T. Hoskins, M. Youngquist, M. Dietrich, A. Christman, L.Wengler, A. Burrow, and T. Keesling for assistance in animal care and collection. We appreciate C. Williamson and E. Overholt for providing materials and lab space for culturing Bd. Funding for this research was provided by Miami University.

